# Efficient multi-lineage cardiovascular differentiation of human pluripotent stem cells in animal serum-free conditions

**DOI:** 10.64898/2026.01.28.702392

**Authors:** Nguyen T N Vo, Kelvin Chung, Aishah Nasir, Davor Pavlovic, Chris Denning

## Abstract

Human induced pluripotent stem cell (hiPSC) technologies offer human-relevant cardiac models for biomedical applications. However, workflows for differentiation of cardiac stromal cells and fabrication of engineered heart tissue (EHT) commonly rely on animal serum, contrary to growing policy demands to reduce use of these products. Applying marker analysis via COL1A, DDR2 and GATA4 for cardiac fibroblasts or CD31, CD34 and CD144 for endothelial cells, we tailored Panexin, a defined serum substitute, to support high efficiency differentiation of cardiac stromal lineages to 85% purity without additional purification steps. We evaluated fabrication of EHTs using hiPSC-cardiomyocytes only (monoculture) or further combined with cardiac fibroblasts and endothelial cells (triculture; 70%:15%:15%, respectively). Panexin poorly supported fabrication and contractility of EHTs, a finding unaltered by modulating spontaneous cardiac myofibroblast activation via TGFβ inhibition. In contrast, human serum enabled fabrication of mono- and tri-culture EHTs, wherein constructs made without TGFβ signalling inhibition delivered the strongest contractile forces (up to 0.25 mN) and exceeded comparator tissues engineered using animal serum. Our data show that iterative evaluation of serum substitutes, human serum, cell combinations and signalling pathway modulators can mitigate use of animal serum for functional EHT generation, aligning with the UK government’s roadmap for alternative methods.

## INTRODUCTION

Human induced pluripotent stem cells (hiPSCs) provide a renewable source of human cardiac cell types for disease modelling, drug discovery and safety pharmacology, and emerging regenerative applications. Although hiPSC-derived cardiomyocytes (CMs) are widely used, co-culture with cardiac stromal cells such as cardiac fibroblasts (CFs) and endothelial cells (ECs) often improves tissue organisation, contractile performance and maturation, and can better capture disease phenotypes *in vitro* ^1–4^.

Despite the widespread use of animal serum (e.g. fetal bovine serum, FBS; bovine serum albumin, BSA) in cell culture since the 1950s, its inclusion in differentiation protocols raises several concerns^5^. These include ethical issues, physiological relevance and the undefined nature of animal serum. Introduction of non-human growth factors or cytokines may activate unintended pathways, hindering efforts to identify specific factors responsible for biological effects hence impeding mechanistic studies and protocol optimisation^6–9^. Animal serum can introduce contamination risks (e.g., viruses, prions) and immunogenic responses, complicating therapeutic and regulatory compliance^5,7,10^. These limitations obstruct scaling for industrial applications and regulatory approval. Consequently, there is a shift towards non-animal methods (NAMs), illustrated by the UK government’s recent policy document, “Replacing animals in science: A strategy to support the development, validation and uptake of alternative methods”^11^.

To address these limitations, chemically defined substitutions have been developed. This includes Panexin NTA pharma-grade, designed to support adherent cell culture using defined and human-compatible components for improved reproducibility and regulatory compliance. In addition to serum-free substitutes, human serum is a biologically relevant alternative that reflects human physiology, although challenges of variability and availability remain. Of note, engineered heart tissue (EHT) platforms typically rely on animal serum during both fabrication and long-term culture to support compaction, tissue remodelling and contractile maturation.

In this study, we tested two complementary strategies to reduce reliance on animal serum across a multi-lineage cardiac modelling workflow. First, we evaluated Panexin for use in differentiating hiPSC-derived CFs and ECs without the need for further purification or enrichment. Second, we examined whether Panexin or human serum could replace animal serum during EHT fabrication from monocultures of hiPSC-CMs (cardiomyocyte-only) and tricultures (tri-lineage) of hiPSC-CMs, CFs and ECs. This included exploring the impact of TGFβ pathway modulation of spontaneous cardiac myofibroblast activation on EHT function. The results below show how we reached these conclusions, ending with a summary infographic of idealised guidelines for others in the field.

## RESULTS

### PANEXIN enables serum-free differentiation of cardiac fibroblasts at high purity

We had previously optimised cardiomyocyte differentiation of two in-house hiPSC lines, REBL-PAT and AT1^12^, providing the grounding to develop approaches to produce cardiac stromal cell from the same lines. For CFs, we adapted a differentiation protocol from Zhang et al^13^. Fetal calf serum (FCS) in the Promocell fibroblast growth medium (FGM) was replaced by Panexin at an equivalent volume, generating a serum-free version of FGM (figure 1.A, and supplementary figure S1). We then trialled differentiation using seeding densities from 20,000 to 30,000 cell per cm^2^. Timelines for differentiation were adapted, with cells replated at day 14 post-differentiation for onward culture in serum-free FGM supplemented with 10 ng/ml FGF2 and 10 µM SB431542.

**Figure 1.**
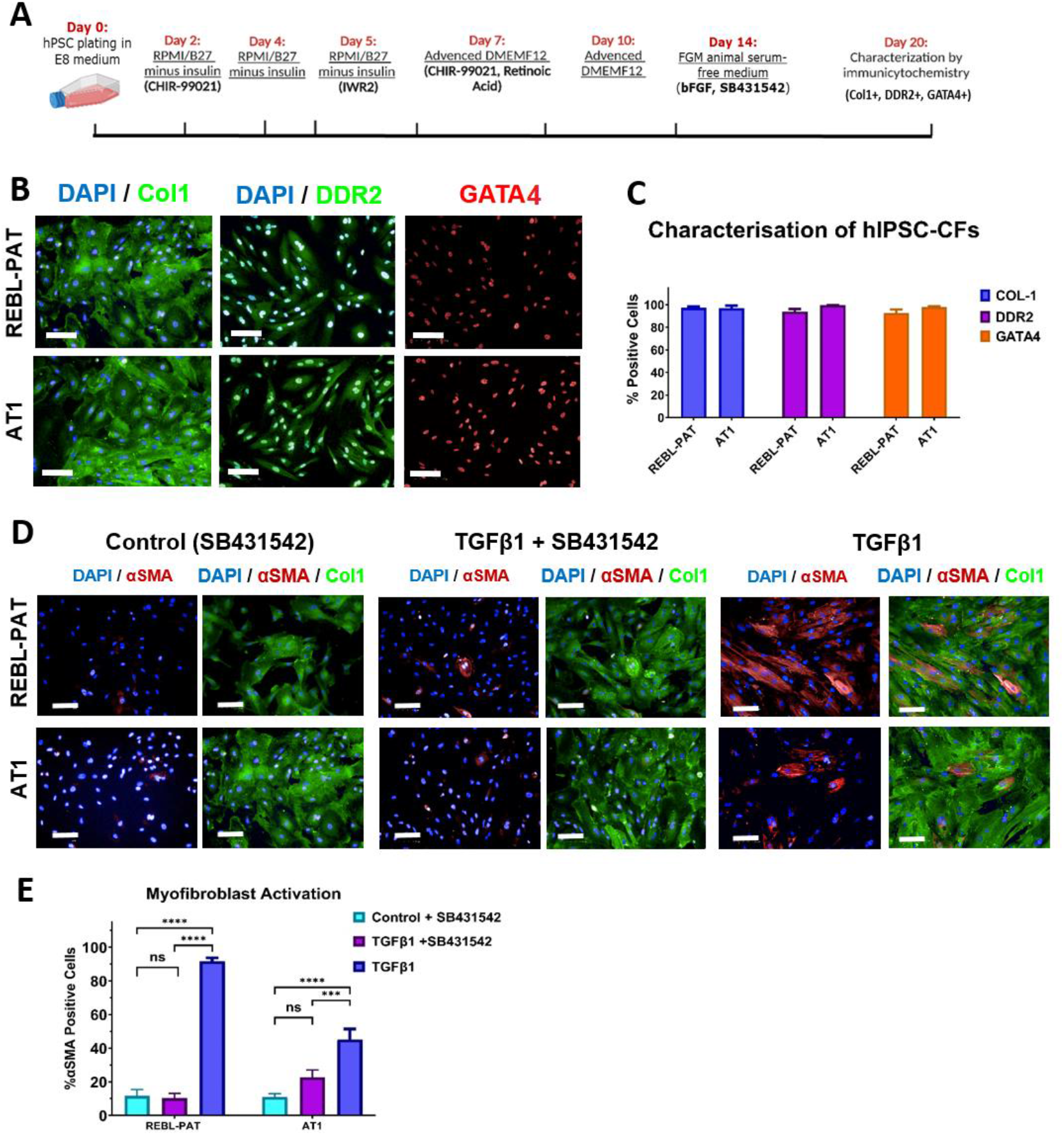
Serum-free differentiation of hiPSC-derived cardiac fibroblasts using Panexin. (A) Schematic of the modified protocol. (B) Representative immunostaining illustrating the expression of CF-specific markers, including COL1A & DDR2 (green) and GATA4 (red), scale bar: 100 µm. (C) Quantification of CF purity, presented mean ± SEM, N = 3 biological replicates. (D) Myofibroblast activation assay under TGF-β stimulation with and without SB431542. Nuclei (DAPI in blue), COL1A (green), α-SMA (red), scale bar = 100 µm. (E) Quantification of αSMA-positive cells, presenting mean ± SEM, N = 3 biological replicates. Two-way ANOVA with Tukey’s multiple tests; ns, not significant.

Analysis by immunostaining at day 20 post-differentiation revealed high proportions of COL1+ and DDR2+ fibroblasts co-expressing the cardiac-associated marker GATA4 (figure 1.B). Across biological replicates, more than 90% of cells were marker-positive in both lines (figure 1.C), comparable to efficiencies obtained using the serum-based protocol (supplementary figure S1). Taken together, these data show that modified differentiation conditions using Panexin supports high efficiency serum-free production of CFs, which display robust expression of both fibroblast and cardiac-associated markers.

### TGFβ-driven myofibroblast activation is preserved under serum-free conditions and suppressed by SB431542

To assess functional responsiveness, hiPSC-CFs generated under serum-free conditions were treated with TGFβ1 in the presence or absence of the TGFβ receptor inhibitor, SB431542. TGFβ1 induced a pronounced increase in alpha smooth muscle actin (αSMA) expression accompanied by a myofibroblast-like morphology, whereas co-treatment with SB431542 markedly reduced αSMA induction, consistent with inhibition of TGFβ signalling (figure 1.D).

In the REBL-PAT line, TGFβ1 stimulation resulted in 91.7% αSMA-positive cells, while vehicle-treated cells and TGFβ1-treated cells in the presence of SB431542 showed low αSMA expression (<20%; figure 1.E). A similar response was observed in the AT1 line, with 45.2% αSMA-positive cells following TGFβ1 treatment compared with 11% in vehicle controls and 22.7% in cells treated with both TGFβ1 and SB431542. Overall, hiPSC-CFs generated under Panexin-based serum-free conditions retain expected responsiveness to TGFβ signalling. The modulation of myofibroblast activation closely parallels that observed in serum-based differentiation protocols (supplementary figure S1), supporting the functional equivalence of CFs derived using Panexin.

### Panexin supports serum-free endothelial differentiation without purification

Endothelial differentiation was adapted from our previously published protocol^12^ by replacing bovine brain extract (BBE) and fetal bovine serum (FBS) in the commercial Lonza EGM™ Endothelial Cell Growth Medium BulletKit™ with an equivalent volume of Panexin to generate a serum-free formulation of EGM (figure 2.A). Optimisation of cell seeding density, ranging from 20,000 – 40,000 cells per cm^2^ was required for both serum-free and serum-based conditions (supplementary figure S2).

**Figure 2.**
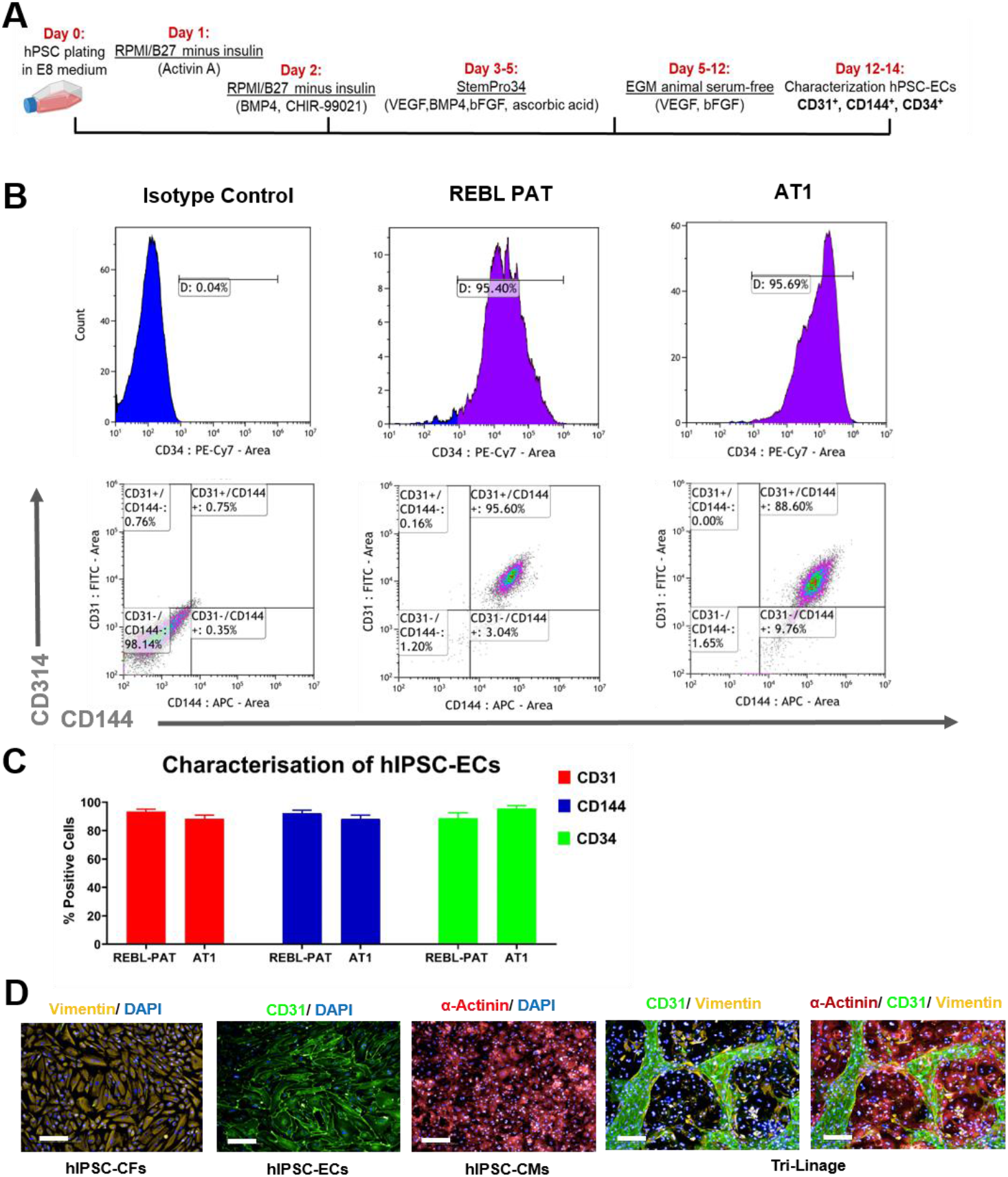
Serum-free differentiation of hiPSC-derived endothelial cells using Panexin. (A) Schematic of the modified protocol. (B) Representative flow cytometry for EC-specific markers CD31, CD144 (VE-Cadherin) and CD34 in REBL-PAT and AT1. (C) Quantification of EC purity generated by the serum-free protocol, presented mean ± SEM, N = 3 biological replicates. (D) Representative images illustrate the angiogenesis establishment in the tri-culture of hiPSC-ECs, hiPSC-CFs and hiPSC-CMs, Vimentin (yellow), CD31 (green) and α-actinin (red), scale bar 200 µm.

Following replating and maturation, flow cytometry on day 13 showed greater than 85% positive cells for CD31, CD144 and CD34 without the need for magnetic- or FACS-based purification (figure 2.B - C). Preserved endothelial functionality was illustrated in triculture monolayers of CMs, CFs and ECs, wherein vascular-like networks were evident after 7 days of co-culturing (figure 2.D).

### Panexin does not support functional contractile monoculture hiPSC-CM EHTs

Strip-format fibrin-based EHTs^14^ are widely used to assess contractile responses of human and animal cardiac tissues in 3D format. As this protocol requires FCS (10% v/v) during fabrication and horse serum (10% v/v) during maintenance culture, we evaluated whether Panexin could replace animal serum during fibrin-based EHT formation and long-term culture (supplementary figure S3.A & B).

EHTs were fabricated using monocultures of hiPSC-CMs, also termed CM-only EHTs, and maintained in medium supplemented with Panexin at 10%, 1%, or 0.5% (v/v) (figure 3A). Tissue morphology and contractile activity were monitored over a 21-day period using serial imaging and video-based force analysis (supplementary figure S3.C-D).

**Figure 3.**
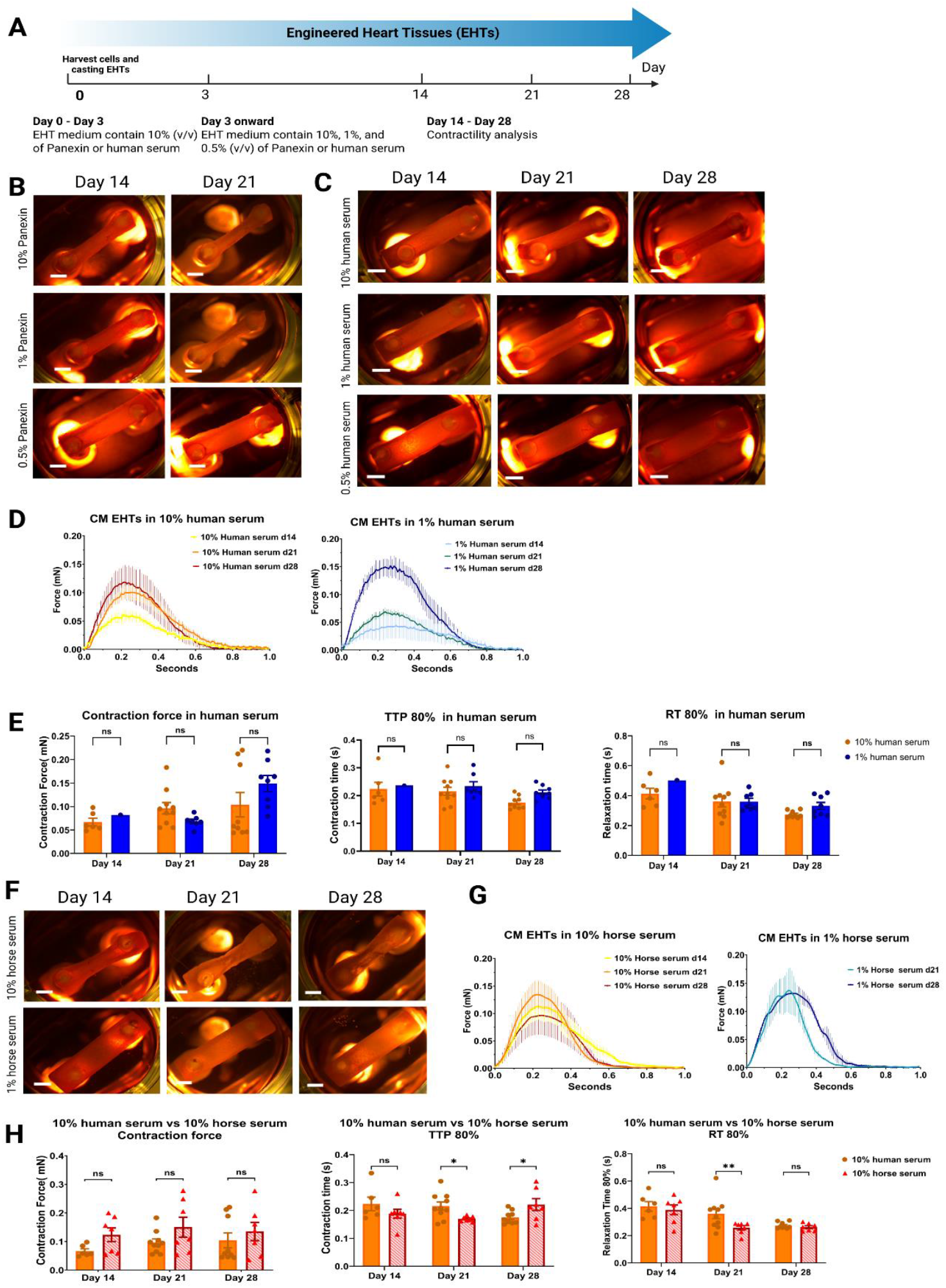
Serum replacement in monoculture EHTs. (A) Workflow schematic. (B) Representative morphology over time in Panexin conditions. (C) Representative morphology over time in human serum conditions, scale bar 2mm. (D) Contraction traces of CM-only EHTs in 10% and 1% v/v human serum. (E) Comparison of contraction force, contraction time, and relaxation time between low (1% v/v) and high (10% v/v) human serum concentrations. (F) Representative morphology in CM-only EHTs in horse serum, animal serum control. (G) Contraction traces of CM-only EHTs cultured with horse serum at high (10% v/v) and low (1% v/v) concentrations. (H) Comparison of contraction force, contraction time, and relaxation time between human serum and horse serum at 10% v/v. Data are presented as mean ± SEM (n = 3 biological replicates, 2-4 technical replicates per condition). Two-way ANOVA with Tukey’s multiple comparison test; ns, not significant, * p ≤ 0.05, ** p ≤ 0.01, and *** p ≤ 0.001.

Morphologically, EHTs cultured with 10% Panexin exhibited defined borders and progressive remodelling from an initial rectangular geometry to a characteristic dumbbell shape (figure 3.B), similar to tissues generated using animal serum (figure 3.F). Despite preserved structural integrity, these EHTs failed to exhibit spontaneous beating or contractile responses upon electrical pacing at 1 Hz. Reducing Panexin from 10% to 1% or 0.5% (v/v) compromised EHT structural integrity, remodelling, and function, with no detectable contractile activity, either in spontaneous or paced conditions. Tissue formation and stability were further reduced under serum-free (0%) conditions, where low levels of cell attachment to the silicon posts were observed, and remaining tissues frequently detached or failed to exhibit contractile responses during maturation (supplementary figure S3.C). Together, these findings indicate that, while Panexin supports morphological remodelling of CM-only EHTs at higher concentrations, it is insufficient to sustain functional contractility under all tested concentrations.

### Human serum supports functional EHT fabrication and contractility

Given the limitations of Panexin in supporting functional CM-only EHTs, we next investigated whether human serum could replace animal serum under otherwise identical fabrication and culture conditions. EHTs generated in the presence of 10% or 1% human serum displayed well-defined borders and robust remodelling over time, whereas tissues formed using 0.5% human serum failed to support EHT construct structure and contractility (figure 3.C).

Functional assessment revealed that the monoculture EHTs cultured with either 1% or 10% human serum generated contractile forces of approximately 0.1 mN by day 21 (figure 3.D & E). This was broadly comparable to tissues produced using animal sera, although in some metrics, such as consistency of contraction force (figure 3.F & G), 1% horse serum performed less robustly than 1% human serum. Contractile parameters, including contraction time and relaxation time, were comparable between human and animal serum at 10% v/v concentration (figure 3.H). These results demonstrate that human serum can effectively support both morphological maturation and functional contractility of CM-only EHTs, providing a human-relevant alternative to animal serum.

### Omission of TGFβ inhibition enhances triculture EHTs contractility

Cardiac fibroblasts and endothelial cells can confer a positive impact on cardiomyocyte function and maturation through extracellular matrix deposition and paracrine signalling. We therefore investigated whether incorporation of cardiac stromal cells would influence morphological, structural, or functional properties of EHTs under animal serum-free conditions. Triculture EHTs were generated using a defined cellular composition of 70% hiPSC-CMs, 15% hiPSC-CFs, and 15% hiPSC-ECs.

To support all three cell types within a single construct, a universal EHT culture medium was used for both mono- and tri-culture tissues throughout this study. This medium consisted of DMEM supplemented with 1% penicillin/streptomycin, 0.1% insulin, 0.1% aprotinin, and either Panexin or serum (human or horse) at 10% or 1% (v/v), with the remaining volume made up by DMEM. This formulation was selected to maintain compatibility across CMs, CFs, and ECs while enabling direct comparison of serum and serum-replacement conditions.

Based on poor performance in previous experiments, Panexin and human serum concentrations of 0% and 0.5% were excluded, as was 1% horse serum. Vascular endothelial growth factor (VEGF; 30 ng/mL) and fibroblast growth factor 2 (FGF2; 10 ng/mL) were added into the universal EHT medium culture to support ECs and CFs within the triculture constructs. To modulate spontaneous myofibroblast activation, EHTs were cultured in the presence or absence of the TGFβ receptor inhibitor SB431542 (10 µM) (figure 4.A; supplementary figure S4).

**Figure 4.**
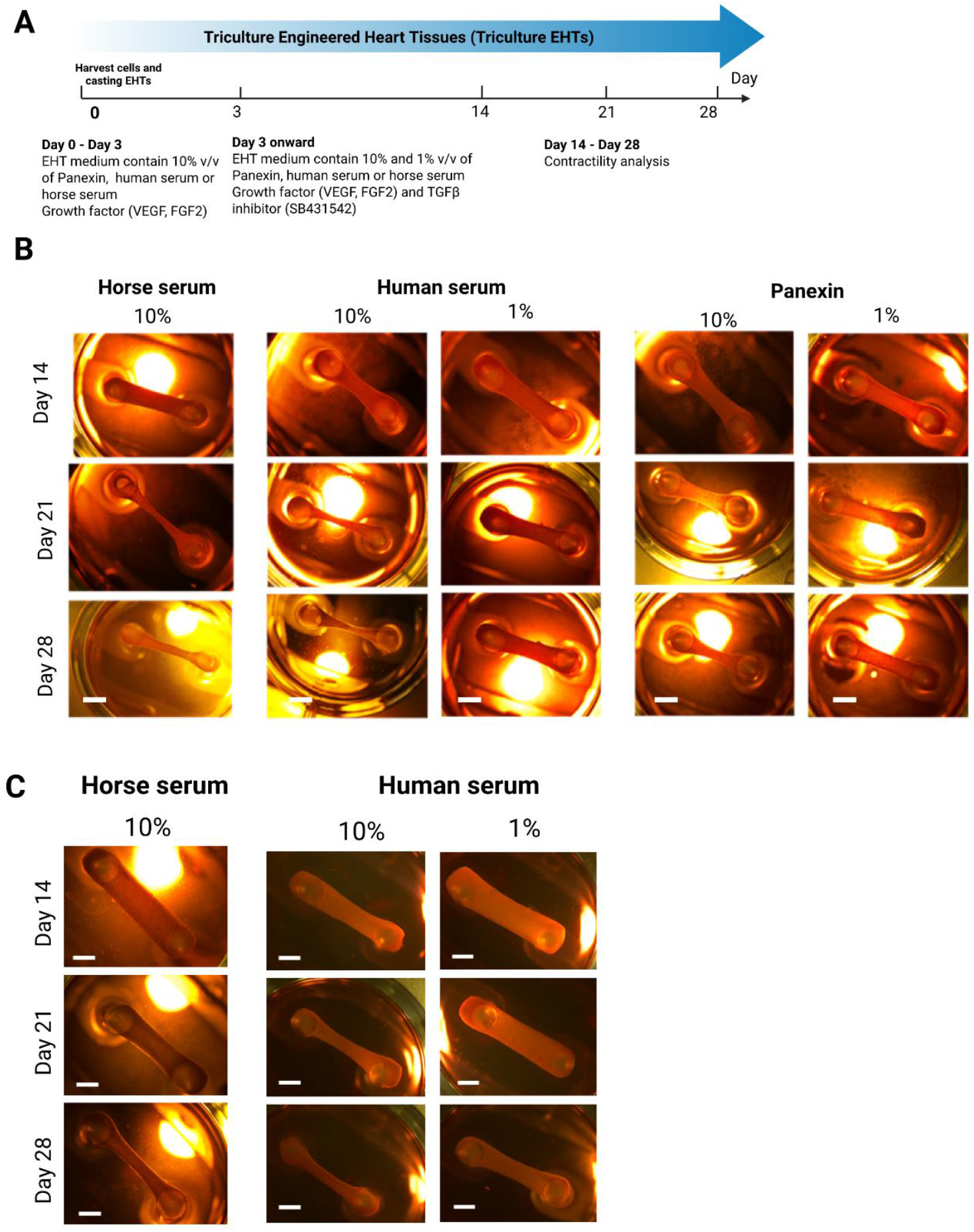
Triculture EHT morphology under serum replacement. (A) Workflow schematic for triculture EHTs. Representative morphology of EHTs over time across serum and serum-replacement conditions with SB431542 (B) and without SB431542 (C), scale bar 2 mm.

Across all tested conditions (1% or 10% serum or Panexin, with or without SB431542), triculture EHTs exhibited well-defined boundaries and pronounced remodelling by day 14. This further increased by day 21 (figure 4.B-C), indicating a positive structural contribution from the cardiac stromal populations. However, marked differences in contractile performance were observed depending on SB431542 treatment (figure 5).

**Figure 5.**
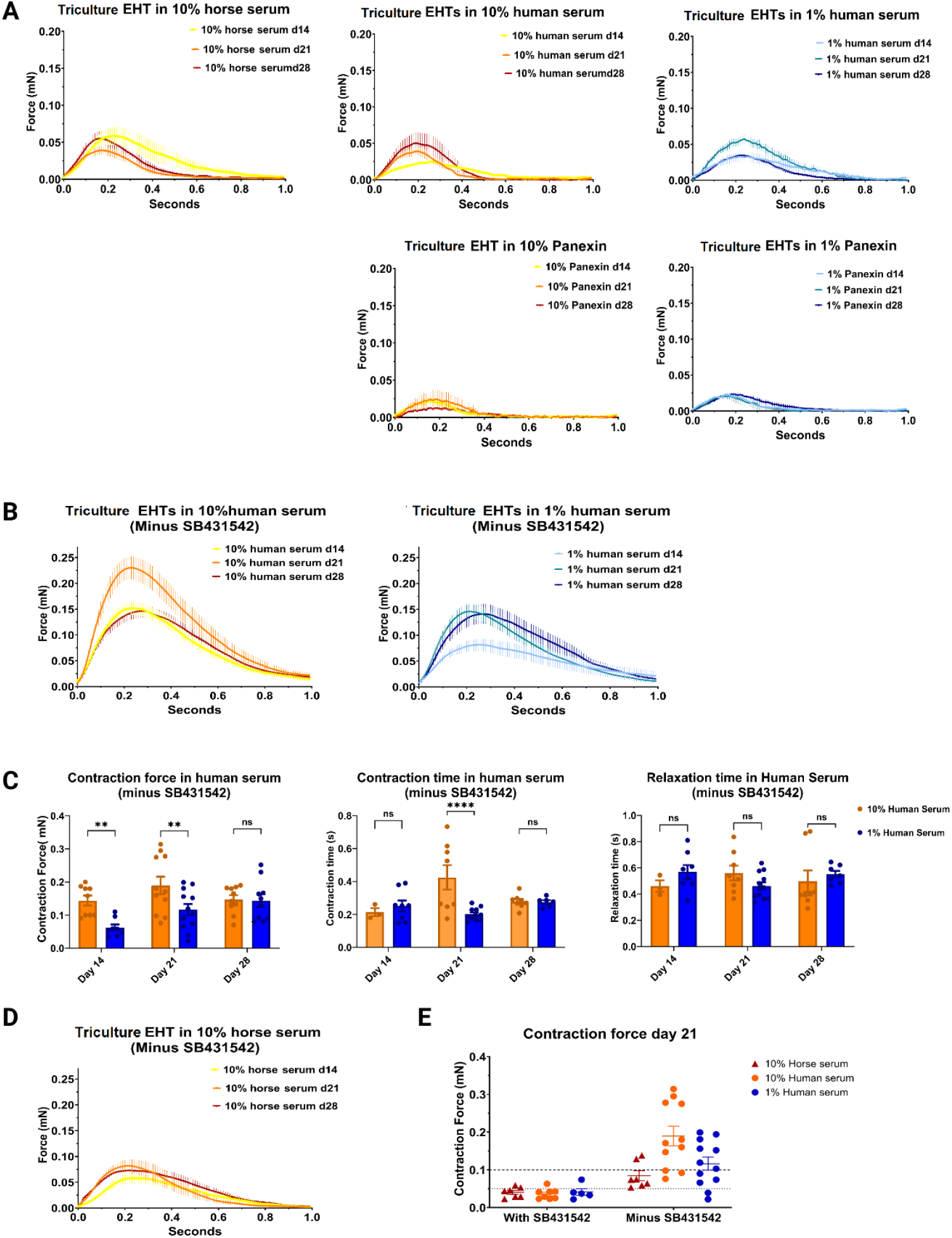
Contractile analysis of triculture EHTs and effect of SB431542. (A) Contraction traces of triculture EHTs across serum and serum-replacement in the presence of SB431542. (B, C) Contraction traces and contractile metrics (contraction force, contraction time, and relaxation time) of triculture EHTs cultured with 10% and 1% v/v human serum in absence of SB431542, data representing means ± SEM, n=3 biological replicates, 2-4 technical EHT replicates. Two-way ANOVA with Tukey’s multiple comparison tests. (D) Contraction traces of triculture EHTs cultured with 10% horse serum in the absence of SB431542. (E) Direct comparison of contraction force across conditions at late time points at d21 post-fabrication of triculture EHTs in all serum conditions in the presence and absence of SB431542. Data representing means ± SEM, n=3 biological replicates, 2-4 technical EHT replicates, two-way ANOVA with Sidak’s multiple comparison test; ns, not significant, * p ≤ 0.05, ** p ≤ 0.01, and *** p ≤ 0.001.

In the presence of SB431542, contraction forces were substantially reduced across all conditions, reaching only 0.02 mN in Panexin-supported EHTs, and around 0.05 mN in human serum-supported and horse serum-supported EHTs (figure 5.A). In contrast, omission of SB431542 restored contractile performance in both 1% and 10% human serum conditions, with forces attaining or exceeding 0.1–0.15 mN by day 21 (figure 5.B-C). These values were comparable to, or greater than, those observed in monoculture EHTs (figure 4).

Omission of SB431542 in triculture EHTs cultured with horse serum also showed an increase in contractile force (figure 5.D). Direct comparison demonstrated that triculture EHTs cultured with 1% human serum closely aligned with the contractile performance observed in the 10% v/v animal serum or human serum conditions from day 21 onward (figure 5.E). This comparative analysis across all serum conditions revealed a pronounced negative impact of SB431542 on contraction force. Despite the positive contribution of cardiac stromal cells, triculture EHTs cultured with Panexin did not show significant improvement in contractile force (∼0.04 mN) following SB431542 removal (supplement figure S5), which remained insufficient for functional studies requiring robust contractile output.

## DISCUSSION

As summarised in Figure 6, this study provides a guide on the conditions required to negate use of animal serum in differentiating cardiovascular lineages from hiPSC and then fabricating these into mono- or tri-culture engineered heart tissues.

**Figure 6.**
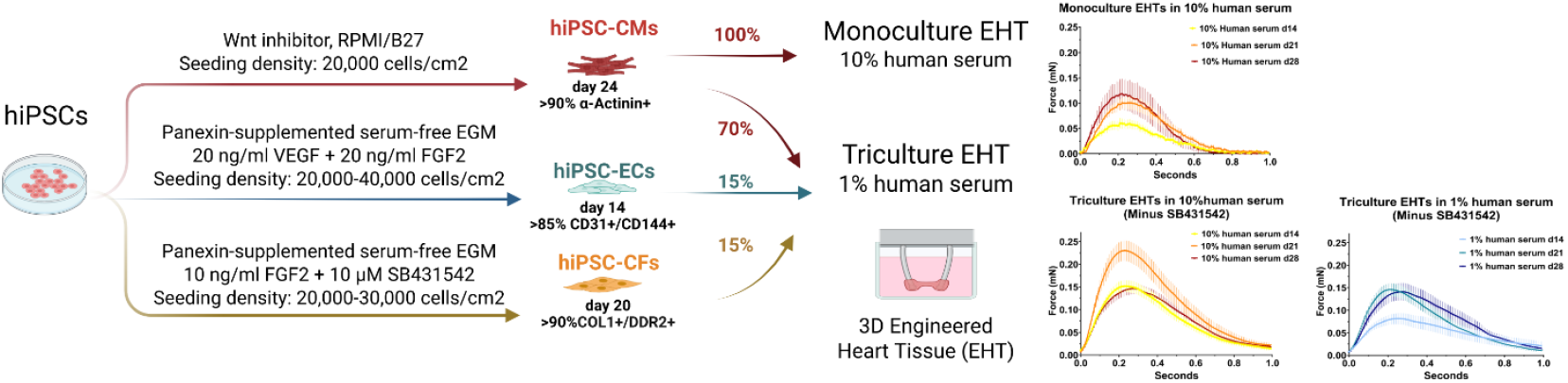
Summary of serum-free stromal differentiation and optimised serum conditions for engineered heart tissue assembly. Schematic overview of the workflow developed in this study. Human iPSCs were differentiated into cardiomyocytes (CMs), endothelial cells (ECs), and cardiac fibroblasts (CFs) using serum-free protocols, wherein fetal bovine serum was replaced by Panexin for stromal lineages. For engineered heart tissue (EHT) fabrication, 10% v/v human serum efficiently supported morphological remodelling and functional contractility in CM-only EHTs. Triculture EHTs assembled from CMs (70%), ECs (15%), and CFs (15%) showed enhanced structural organisation and contractile performance under high (10% v/v) and low (1% v/v) human serum conditions in the absence of prolonged TGFβ inhibition by SB431542. Representative contraction traces illustrate functional differences between monoculture and triculture EHTs under optimised conditions.

These advances in serum-free differentiation protocols for cardiac stromal cells build on our previous work^15^ by demonstrating scalable, purification-free generation of both CFs and ECs using Panexin. While several serum-free and animal component-free media have been reported for cardiac differentiation and three-dimensional cardiac microtissues ^16–18^, these approaches often require additional purification steps to achieve high-purity stromal populations. In contrast, Panexin supported robust differentiation of CFs and ECs without sorting, reducing hands-on time and improving scalability.

However, Panexin did not support functional contractile maturation in fibrin-based CM-only EHTs, indicating that defined serum substitutes suitable for two-dimensional stromal differentiation may lack key components required for three-dimensional tissue assembly and electromechanical coupling. Although incorporation of cardiac stromal cells improved structural remodelling in triculture EHTs, contractile performance in Panexin-based conditions remained inconsistent and insufficient compared with animal serum controls^14,19^. This is likely because Panexin is less complex, contains fewer nutrients and has no growth factors with only 2% v/v human-derived components. This notion was confirmed when we attempted to fabricate EHTs in the absence of serum, wherein a low level of cell attachment to the silicon posts during and post the casting process was observed. During the culturing duration, the remaining tissues of no-serum EHTs detached from the posts or showed no signs of contractility response.

Human serum is a functional alternative for EHT fabrication and maintenance. At low concentration (1%), human serum consistently supported EHT formation and contractile force comparable to animal-serum controls, while potentially reducing non-physiological high-serum effects. Notably, low concentrations of human serum may minimise hypertrophic and fibrotic responses compared to higher serum concentrations, reducing experimental artefacts and enhancing applicability for disease using hiPSCs models. This study showcases the utility of human serum in supporting cardiac tissue engineering while maintaining physiological relevance and also align with recent serum-free studies ^18,20^.

Although challenges remain to using human serum, such as batch-to-batch variability, limited scalability, and logistical constraints, particularly hormone cycles for female donors^18,21^, the closer physiological relevance and the ability to use low concentrations make it an attractive interim option for NAM-aligned cardiac tissue models. Moreover, human serum is subject to extensive donor screening and quality control procedures, including viral testing (e.g., HIV, hepatitis B and C), sterility, and endotoxin assessment at the blood bank and manufacturing stages^7,22–24^. Such standardised screening and processing steps can mitigate safety risks and support translational use, particularly when combined with low serum concentrations and defined culture conditions.

A key finding with practical implications is the reduction in triculture EHT contractile force following prolonged exposure to the TGFβ receptor inhibitor SB431542. SB431542 is widely used to limit myofibroblast activation and has been shown to protect cardiomyocytes from contractile dysfunction driven by excessive TGFβ signalling under fibrotic or pathological stress conditions ^13,18,25^. In contrast, our data indicate that sustained TGFβ pathway inhibition compromises contractile performance under non-fibrotic baseline conditions. A plausible explanation is disruption of essential extracellular matrix (ECM) remodelling, as SB431542 inhibits fibroblast-mediated collagen gel contraction, a process required for tissue compaction and mechanical conditioning^26–28^. Importantly, this observation does not contradict reports in which SB431542 improved or preserved contractility in human 3D cardiac models, as those studies typically employed short-term exposure during or after pathological stimulation^29–31^.

In contrast, prolonged exposure (up to 28 days) in our healthy triculture EHTs may reveal context-dependent effects on physiological ECM dynamics. These findings highlight the importance of duration- and context-specific optimisation of TGFβ inhibition in human three-dimensional cardiac constructs^32^. Future work should therefore explore strategies such as timed dosing or reduced inhibitor concentrations to balance suppression of excessive fibrosis with preservation of necessary matrix mechanics. Our finding in using low human serum concentrations (1% v/v) to sufficiently support functional triculture EHTs further underscores the need to interpret TGFβ modulation within defined experimental frameworks.

Overall, our findings support a stepwise transition away from animal serum by: (i) using Panexin for scalable, purification-free cardiac stromal lineage differentiation; (ii) using low-concentrations of human serum for functional EHT generation; and (iii) re-optimising anti-fibrotic interventions such as SB431542 within triculture tissues. Together, these advances align with 3Rs principles and strengthen the translational relevance and reproducibility of hiPSC-based cardiac models. Although significant progress has been made, further research is needed to refine protocols for 3D tissue generation, enhance functional maturation, and address practical challenges associated with serum replacement. These efforts will pave the way for the development of reproducible, ethical, and clinically relevant cardiac models, advancing both basic research and translational applications in regenerative medicine.

## METHODS

A complete list of reagents, media, supplements, and their suppliers is provided in Supplementary Table S1

### Monolayer cardiac differentiation of hiPSCs

Routine hiPSC maintenance and cardiac differentiation was performed using a monolayer protocol adapted from previously established methods^12^. Briefly, REBL-PAT and AT1 hiPSC lines (passages 27–40) were maintained in Essential 8™ medium and passaged every 2–3 days at 70–80% confluence using TrypLE Select. Monolayer cardiac differentiation began by seeding hiPSCs on day 0 and transitioning to StemPro-34 with BMP4 and Activin A as specified. WNT modulation was achieved using KY02111 and XAV939, followed by culture in RPMI/B27 (with insulin) from day 7–9 onward. Cardiomyocytes were dissociated at day 15 using TrypLE Express and collected by centrifugation at 300×g for 5 min. Cardiomyocyte purity was quantified using cardiac-specific markers, including α-actinin or cardiac troponin T (cTnT).

### Cardiac fibroblast differentiation

Cardiac fibroblast differentiation was adapted from Zhang et al.^13^ with replacement of FCS by Panexin (PAN Biotech, #P04-95750P). Following WNT activation and inhibition phases, cells were replated and matured in fibroblast growth medium (FGM, PromoCell #C-23130) supplemented with Panexin (replacing fetal calf serum), 10 ng/ml FGF2 and 10 μM SB431542, where indicated. Medium was refreshed every 2 days. The cells were characterized on day 20 post-differentiation using fibroblast markers (*COL1A* and *DDR2*) and a cardiac marker (*GATA4*).

### Endothelial differentiation

Endothelial differentiation was adapted from previously published protocol ^15^ with substitution of FBS and BBE by Panexin. Briefly, after mesoderm induction, cells were treated with BMP4, FGF2 and VEGF and replated at day 7 in endothelial growth medium (EGM, Lonza #CC-3124) supplemented with Panexin, plus 20 ng/ml VEGF and 20 ng/ml FGF2. Medium was refreshed every 2 days. The differentiated cells were characterised at day 13 post-differentiation by flow cytometry for endothelial-specific markers, including CD31 (PECAM1), CD34, and CD144 (VE-Cadherin).

### Myofibroblast activation assay

To induce myofibroblast activation, hiPSC-derived cardiac fibroblasts at day 20 post-differentiation were treated with recombinant human TGFβ1 (10 ng/mL) for 48 hours. Where indicated, cells were co-treated with the TGFβ receptor inhibitor SB431542 (10 µM) to suppress pathway activation. Vehicle-treated cells was also supplemented with SB431542 and served as controls. Following treatment, cells were processed for immunostaining and quantitative analysis of αSMA expression.

### Fabrication of EHTs

Engineered heart tissues (EHTs) were generated following protocols ^23^. Briefly, strip-format fibrin-based EHTs were generated in agarose casting moulds with PDMS racks. A master mix containing 1×10^6^ cells per tissue (CM-only or triculture at 70:15:15 CMs:CFs:ECs) was combined with fibrinogen and thrombin and cast between posts. After polymerisation, tissues were cultured for 3–4 weeks in EHT medium containing either horse serum (control), human serum, or Panexin at indicated concentrations. VEGF (30 ng/ml) and FGF2 (10 ng/ml) were added for triculture tissues, and SB431542 (10 μM) was included or omitted as indicated. The video-optical recording and contraction analysis were conducted by EHT Technologies and Consulting Team Machine Vision (CTMV) software.

### Immunocytochemistry

Cells were fixed in 4% paraformaldehyde, permeabilised with 0.05% Triton X-100, blocked with 4% goat serum, and incubated with primary antibodies overnight at 4°C followed by Alexa Fluor secondary antibodies. Nuclei were stained with DAPI. For quantification and measurement of immunocytochemistry results, images conducted from Operetta^®^ High Content Imaging System (PerkinElmer), were processed and analysed using Harmony^®^ High Content Imaging and Analysis Software (PerkinElmer).

### Flow Cytometry

Cells were fixed in 4% paraformaldehyde, permeabilised with 0.05% Triton X-100, blocked with 1% BSA in PBS. The cell pellet was transferred to flow cytometry tubes or V-bottom 96-well plates and then incubated with antibodies. Flow cytometry was conducted using an ID7000 system or BD FACS CANTO II Cytometer. The resulting data were analysed using Kaluza software (Beckman Coulter), with cell debris removed and isotype controls and single-stain controls used for gating.

## Statistical analysis

All data are presented as mean ± standard error (SEM) from at least three independent experiments unless otherwise stated. Two-group comparisons used two-tailed Student’s t-tests; multiple-group comparisons used one-way or two-way ANOVA with appropriate post hoc tests. Statistical significances were * p ≤ 0.05, ** p ≤ 0.01, and *** p ≤ 0.001.

## Data availability

The datasets generated and/or analysed during the current study are available from the corresponding author upon reasonable request

## ACKNOWLEDGMENTS

The work in C.D.’s lab was supported by Animal Free Research UK, Grant 141. Animal Free Research UK is the UK’s leading non-animal biomedical research charity that exclusively funds and promotes human-relevant research that replaces the use of animals, and by the British Heart Foundation grants SP/F/22/150044 and PG/21/10545.

## Author Contributions

Conceptualization: C. D. and N.T.N.V. Methodology: N.T.N.V and K.C. Investigation: C.D, N.T.N.V and K.C. Formal analysis: N.T.N.V and K.C. Manuscript writing and editing: C.D., N.T.N.V, A.N, and K.C. Resources: C.D. Project supervision: C.D, and D.P.

## Ethics / statement

Human induced pluripotent stem cell lines were generated previously under appropriate ethical approval and informed consent^12^. The study did not involve the collection of new human samples. All experiments were performed in accordance with relevant institutional guidelines and regulations

