## Supplement Figure for "Efficient multi-lineage cardiovascular differentiation of human pluripotent stem cells in animal serum-free conditions"

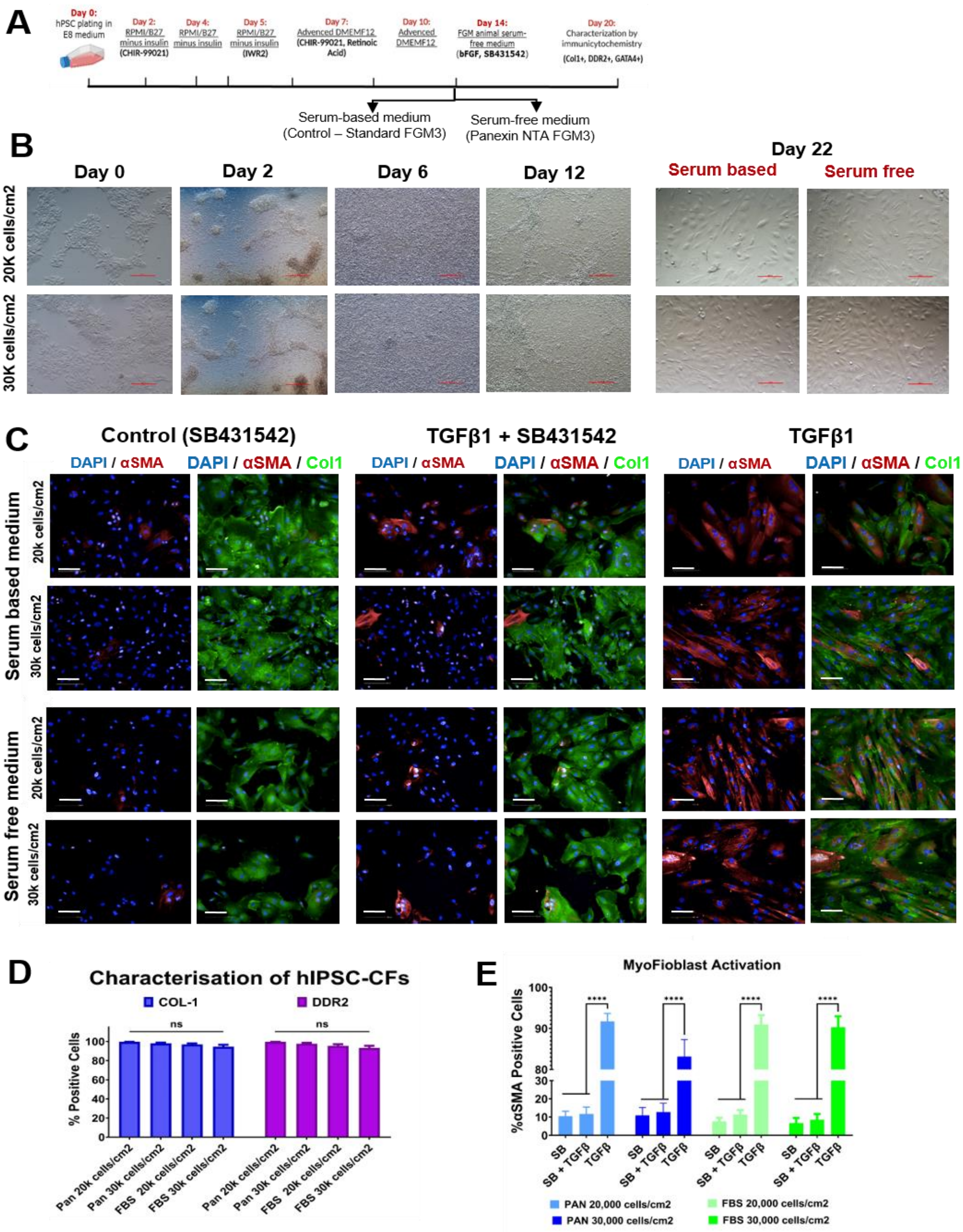

**Supplement Figure S1. Cardiac fibroblast differentiation.**

(A) Schematic of the modified protocol. (B) Cell morphology during cardiac fibroblast differentiation. Scale bar: 500 µm. (C) Immunostaining illustrated myofibroblast transactivation in cardiac fibroblasts differentiated in the serum-based and serum-free protocol with Panexin substitute. Nuclei (DAPI in blue), COL1A (green), α-SMA (red), Scale bar = 100 µm. (D) Quantification of CF differentiation efficiency under various seeding density conditions during the modified serum-free protocol with Panexin. Data presented as mean ± SEM, N = 3 biological replicates. Two-way ANOVA with Tukey's multiple tests; ns, not significant. (E) Quantification of α-SMA positive cells as an indicator of activated myofibroblast induction. Data presented as mean ± SEM, N = 3 biological replicates. Two-way ANOVA with Tukey's multiple tests; ; ns, not significant, * p ≤ 0.05, ** p ≤ 0.01, and *** p ≤ 0.001.

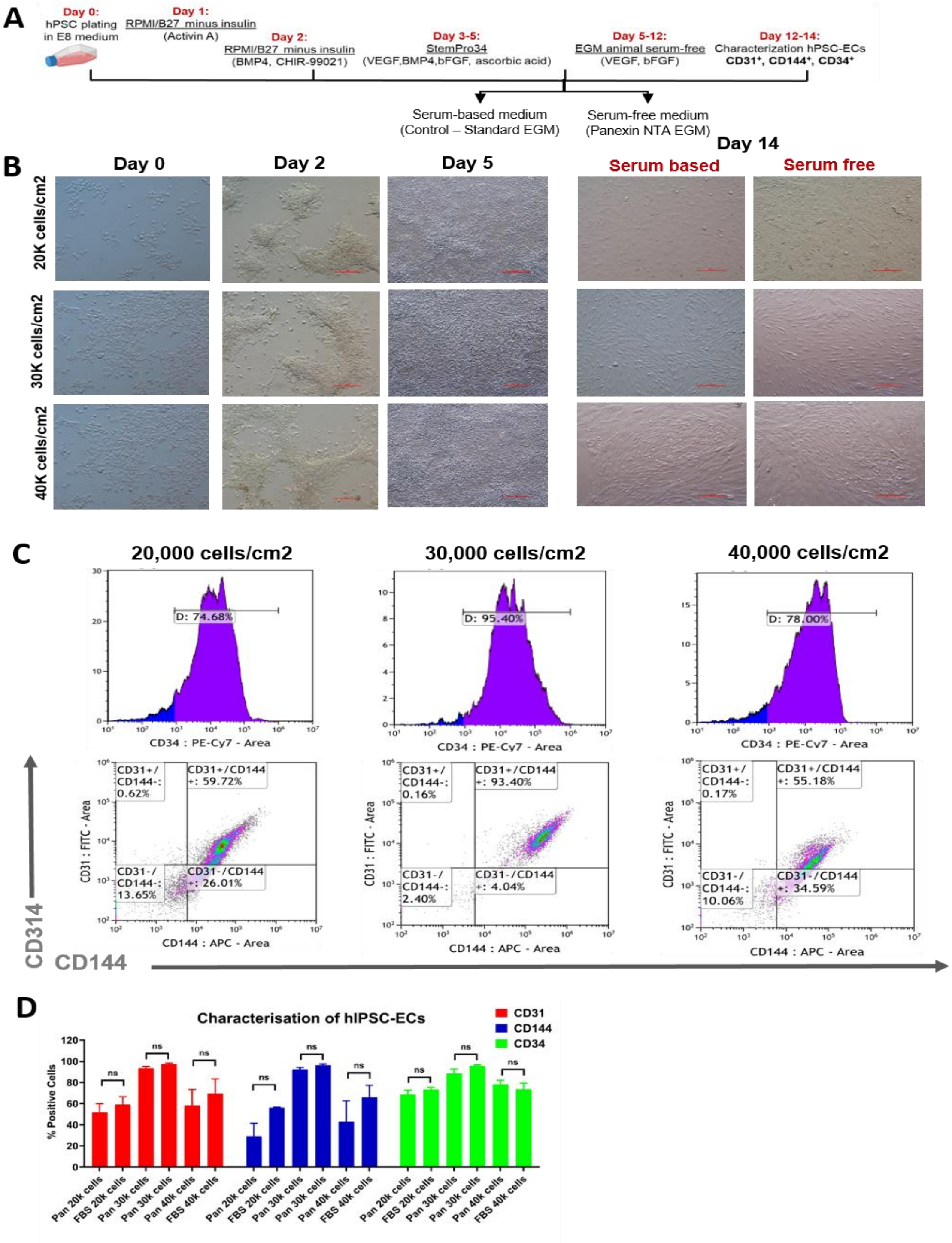


### Supplement Figure 2. Endothelial differentiation.

(A) Schematic of the modified protocol. (B) Cell morphology during endothelial cell differentiation in the serum-based and serum-free protocol with Panexin substitute. Scale bar: 500 µm. (C) Representative flow cytometry plots for EC-specific markers CD31, CD144 (VE-Cadherin) and CD34 on day 14 of differentiation. (D) Quantification EC differentiation efficiency under various seeding density conditions in the serum-free (Panexin) protocol. Data presented as mean ± SEM, N = 3 biological replicates. Two-way ANOVA with Tukey's multiple tests; ns, not significant, * p ≤ 0.05, ** p ≤ 0.01, and *** p ≤ 0.001.

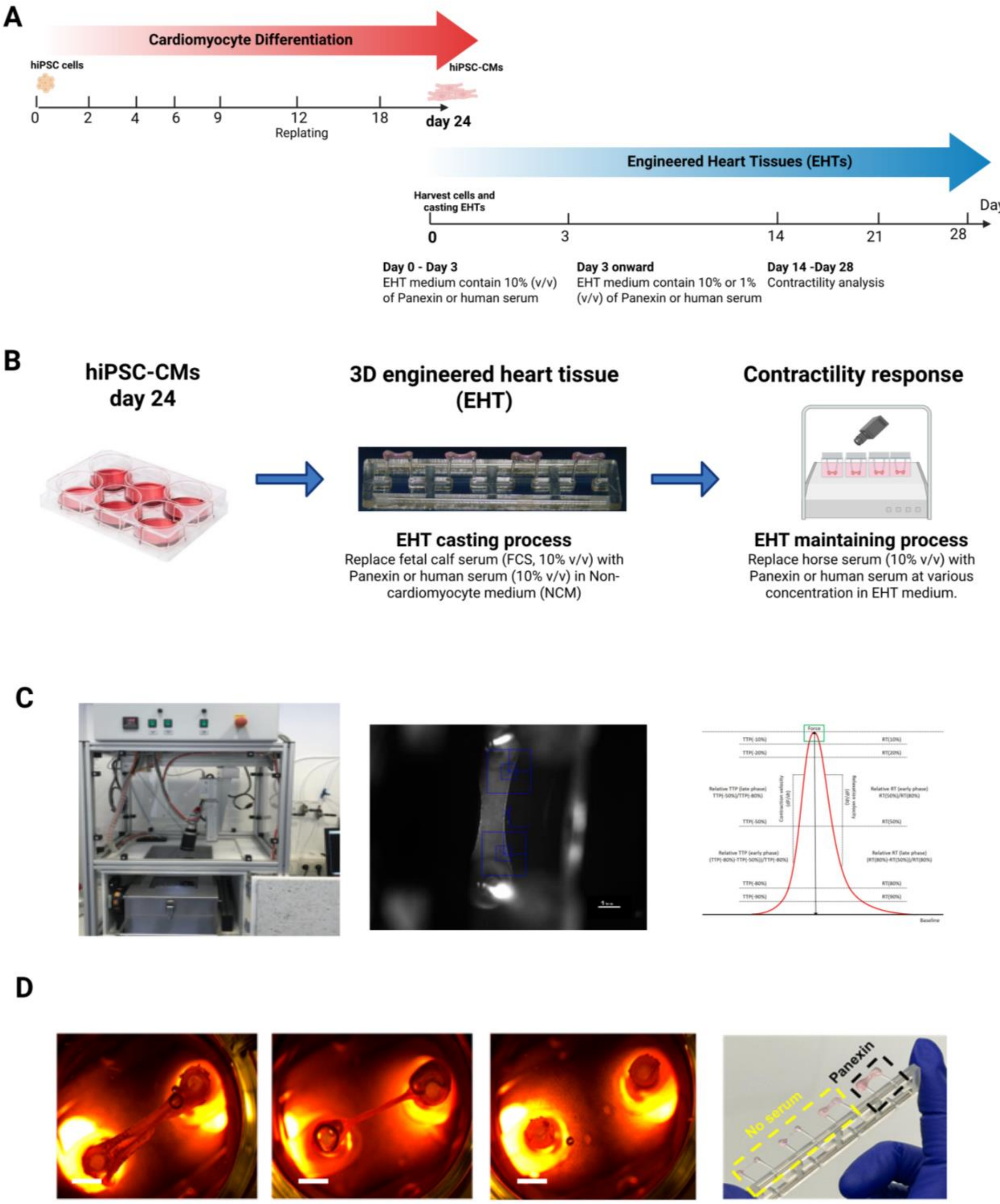


**Supplement figure S3. Evaluation of serum replacement strategies in CM-only EHTs.**
(A) Schematic and timeline for cardiomyocyte differentiation and EHT experimentation. (B) Schematic showing the generation of EHTs using Panexin NTA and human serum to replace the FCS component in the NCM and EHT media. (C) Observations of poor attachment and low yield of tissue constructs under serum-free conditions during tissue fabrication and maintenance. (D) Contraction analysis using the EHT platform and CTMV software, with a diagram explaining the main parameters for contractility analysis, including contraction force, contraction time, and relaxation time.

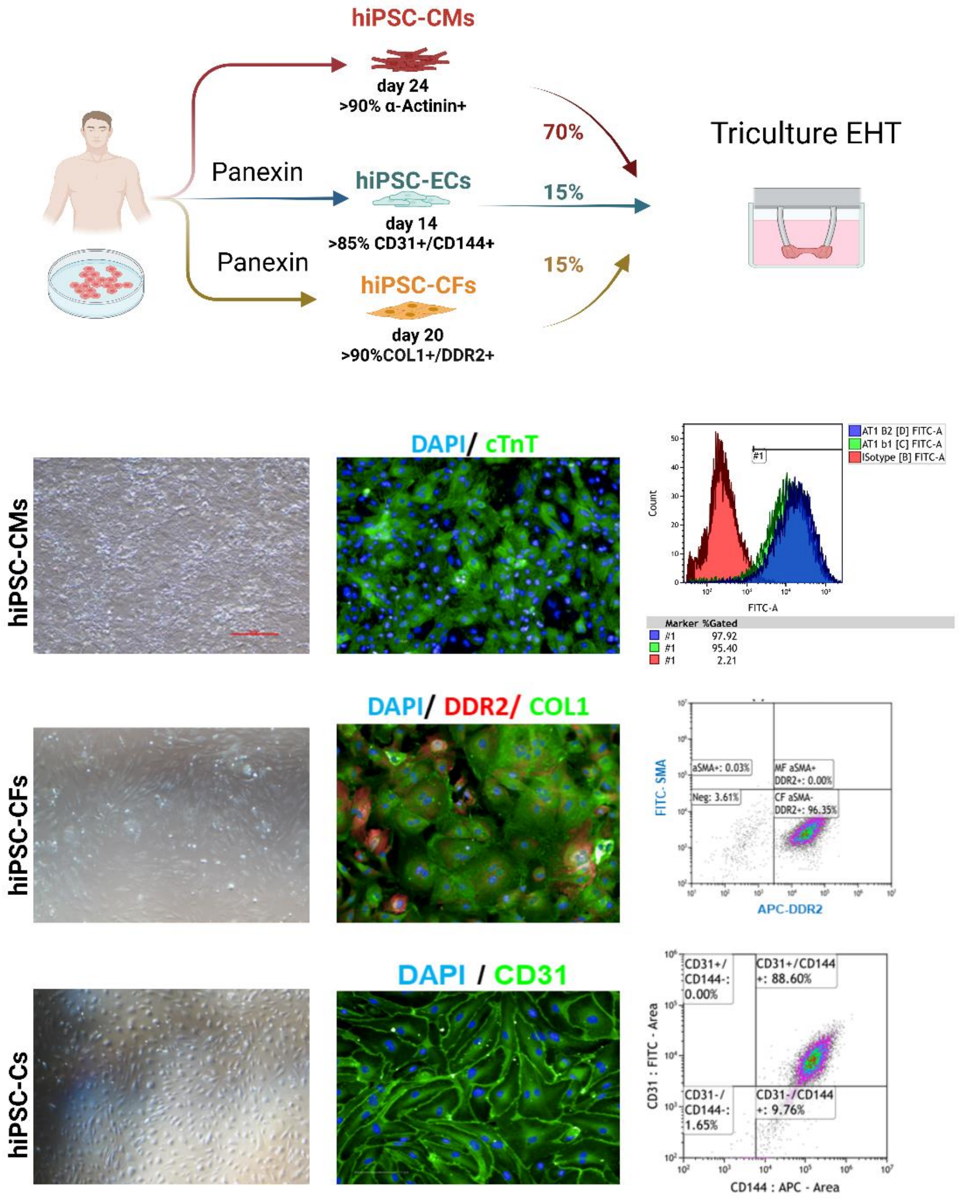


**Supplement Figure 4. Quality control of cell populations used for triculture EHT generation.** Quality control results indicate high purity (>85%) for each cell type, including cardiomyocytes (CMs), cardiac fibroblasts (CFs), and endothelial cells (ECs).

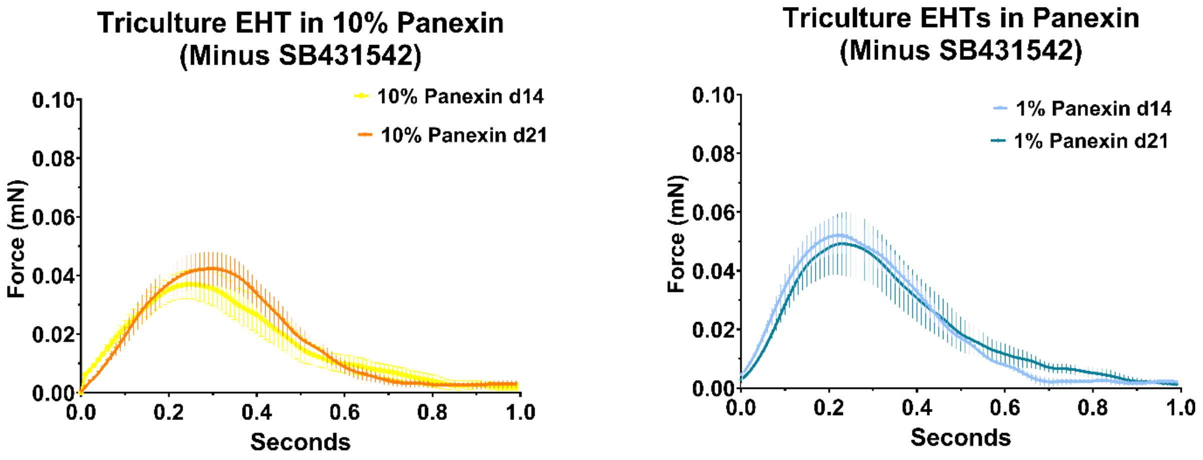


**Supplementary Figure S5. Limited recovery of contractile function in triculture EHTs cultured with Panexin following removal of SB431542.**

Contraction traces of triculture EHTs cultured in Panexin-based medium in the absence of the TGFβ receptor inhibitor SB431542. Despite removal of SB431542 and the presence of cardiac stromal cells, triculture EHTs cultured with Panexin showed only modest contractile forces (~0.05 mN), which remained substantially lower than those observed in human serum conditions. Data presented as mean ± SEM, N = 2 biological replicates, 2-4 technical EHT replicates.

**Supplementary Table S1.** List of reagents, media, and supplements used in this study

| Reagent | Company and Catalogue Number |
|---|---|
| Essential 8™ medium | Thermo Fisher Scientific, A1517001 |
| $Ca^{2+}$ / $Mg^{2+}$ free Phosphate Buffer Saline | PBS, Gibco, 14190-094 |
| TrypLE™ Select (1X) | Thermo Fisher Scientific, 12563029 |
| Y-27632 (Dihydrochloride) | Tocris Bioscience #1254/10 |
| Panexin NTA Pharma Grade | PANBiotech, P04-95700P |
| B-27™ Supplement (50X), minus insulin | ThermoFisher Scientific, A1895601 |
| Advanced DMEM/F-12 | ThermoFisher Scientific, 12634010 |
| Fibroblast Growth Medium 3 | (PromoCell, C-23025) |
| CHIR 99021 | Selleckchem, S2924 |
| IWR-1 | Sigma-Aldrich, I0161 |
| Retinoic Acid | Sigma-Aldrich, R2625-50MG |
| Recombinant Human FGF basic/FGF2/bFGF (146 aa) Protein | R&D, 3718-FB-100 |
| Human TGF-beta 1 Recombinant Protein | PeproTech, 100-21-10UG |
| SB 431542 | Selleckchem, S1067 |
| RPMI 1640 Medium | ThermoFisher Scientific, 11875093 |
| B-27™ Supplement (50X), serum free | ThermoFisher Scientific, 17504044 |
| StemPro™-34 SFM (1X) | ThermoFisher Scientific, 10639011 |
| Vitronectin-N | VN, Lifetech #A14700 |
| Corning® Matrigel® Basement Membrane Matrix, LDEV-free | CORNING, 354234 |
| Human Activin A Recombinant Protein | ThermoFisher Scientific, PHC9564 |
| Recombinant Human BMP-4 Protein | R&D Systems, 314-BP-010/CF |
| KY 02111 | TOCRIS, 4731 |
| XAV 939 | TOCRIS, 3748 |
| Abscobic acid | Sigma, A8960-5G |
| Human VEGF-165 Recombinant Protein | QKine, QK048-0100 |
| Lonza EGM™ Endothelial Cell Growth Medium BulletKit | Lonza, CC-3121 |
| Pan DMEM | PanTechnology, P04-01500 |
| UltraPure™ Agarose | Invitrogen, 15510-027 |
| L-Glutamine (200 mM) | ThermoFisher Scientific, 25030081 |
| Insulin Solution, human recombinant | Sigma, I9278 |
| Newborn Calf Serum | Sigma, N4762 |
| Aprotinin from bovine lung | Sigma, A1153 |
| Penicillin-Streptomycin | Thermo Fisher Scientific, 15070063 |
| Thrombin, Human Plasma | Sigma, 605195 |
| Horse Serum, heat inactivated | Thermo Fisher Scientific, 26050088 |
| Human serum | Sigma, H4522 |
| Fibrinogen from human plasma | Scientific Laboratory Supplies, F4883-500MG |
| Polydimethylsiloxane racks | EHT Technologies C0001 |
| Polytetrafluorethylene spacers | EHT Technologies, C0002 |
| CryoStor® CS10 | STEMcell Technology, 07959 |
| Dimethyl sulfoxide | Sigma, D2650 |
| DMEM, powder, high glucose | Thermo Fisher Scientific, 52100-021 |
| Anti-Collagen I antibody | Abcam, ab34710, working dilution 1:1000 |
| Anti-DDR2 antibody | Abcam, ab63337, working dilution 1:1000 |
| Anti-GATA-4 Antibody (G-4) | Santa Cruz, sc-25310, working dilution 1:1000 |
| Recombinant Anti-Vimentin antibody | Abcam, ab92547, working dilution 1:1000 |

| | |
|---|---|
| Anti-alpha smooth muscle Actin antibody | Abcam, ab7817, working dilution 1:1000 |
| Monoclonal Anti-A-Actinin | Sigma, A7811, working dilution 1:800 |
| Anti-Cardiac Troponin T antibody | Abcam, ab45932, working dilution 1:400 |
| Human CD31/PECAM-1 Antibody | R&D Systems, AF806, working dilution 1:1000 |
| BD Pharmingen™ FITC Mouse Anti-Human CD31 | BD Biosciences, 555445, working dilution 1:20 |
| Alexa Flour 488 Goat anti RabbitCD34 Monoclonal Antibody (4H11), PE-Cyanine7, eBioscience™ | Thermo Fisher Scientific, 25-0349-42, working dilution 1:50 |
| CD144 (VE-cadherin) Monoclonal Antibody (16B1), APC, eBioscience™ | Thermo Fisher Scientific, 17-1449-42, working dilution 1:50 |
| Mouse IgG1, κ Isotype Control | BD Biosciences, 555748 |
| Mouse IGG1K isotype control PECY-7 | Thermo Fisher Scientific, 25-4714-80 |
| Mouse IGG1K isotype control APC | Thermo Fisher Scientific, 17-4714-82 |
| Alexa Flour 488 Goat anti Mouse | Thermo Fisher Scientific, A11029, working dilution 1:1000 |
| Alexa Flour 647 Goat anti Mouse | Thermo Fisher Scientific, A21235, working dilution 1:1000 |
| Alexa Flour 488 Goat anti Rabbit | Thermo Fisher Scientific, A11008, working dilution 1:1000 |
| Donkey anti-Sheep IgG (H+L) Cross-Adsorbed Secondary Antibody AF 488 | Thermo Fisher Scientific, A11015, working dilution 1:1000 |
| Bovine Albumin Fraction V (7.5% solution) | Thermo Fisher Scientific, 15260037 |
| Di-amino phenyl-indole | Sigma, D9542 |
